# Adenosine tips the pathogenic Th1 and Th17 responses in experimental autoimmune uveitis (EAU)

**DOI:** 10.1101/2020.07.06.189183

**Authors:** Deming Sun, Minhee Ko, Hui Shao, Henry J. Kaplan

## Abstract

Various pathological conditions are accompanied by ATP release from the intracellular to the extracellular compartment, where it degrades into adenosine and modulates immune responses. Previous studies concluded that both ATP and its degradation product adenosine are important immune-regulatory molecules; ATP acted as a danger signal that promotes immune responses, but adenosine’s effect was inhibitory. In this study, we show that adenosine plays an important role in balancing Th1 and Th17 pathogenic T cell responses in autoimmune disease. While its effect on Th1 responses is inhibitory, its effect on Th17 responses is enhancing, thereby impacting the balance between Th1 and Th17 responses. Mechanistic studies showed that this effect is mediated via several immune cells, among which γδ T cell activation and dendritic cell differentiation are prominent; adenosine and γδ-mediated immunoregulation synergistically impact each other’s effect. Adenosine augments the activation of γδ T cells, which is an important promoter for Th17 responses and has a strong effect on DC differentiation tipping the balance from generation of DCs that stimulate Th1 responses to those that stimulate Th17 responses. The knowledge acquired in this study should improve our understanding of the immune-regulatory effect of extracellular ATP-adenosine metabolism and improve treatment for autoimmune diseases caused by both Th1 and Th17-type pathogenic T cells.

## Introduction

The purinergic system is an evolutionally selected system modulating immune responses [1, 2]. Under physiological conditions, ATP is contained exclusively within cells; however during tissue damage and inflammation, almost all types of mammalian cells are able to release ATP [2]. Once entering the extracellular space, ATP is hydrolyzed into adenosine diphosphate (ADP), adenosine-5′-monophosphate (AMP), and finally, adenosine in a stepwise manner by ectonucleotidases, including CD73 and CD39 [1, 3-6]. Previous studies have shown that while ATP acts like an endogenously generated TLR ligand capable of augmenting immune responses [7-10], the ATP metabolite adenosine is profoundly anti-inflammatory [11-16]. An increase in extracellular adenosine reduces the local inflammatory response, while removal of endogenous adenosine aggravates tissue dysfunction elicited by injury [17]. Binding of adenosine to its receptors modulates various pathophysiological responses, including immune responses [1, 4-6, 18]. The discovery of the regulatory effect of adenosine on inflammation and immune responses has led to attempts to treat immune dysfunctions by targeting adenosine receptor (AR) signaling [1, 5, 19, 20]. Targeting adenosine receptors and adenosine generation has been successful in treating cancer and neurological diseases [5, 21, 22].

The extrapolation of adenosine as inhibitory was mostly obtained from studies of Th1-type (IFN-γ-producing cells) immune responses, since Th17 responses were discovered only recently. Given the available knowledge that both Th1 and Th17 pathogenic T cells contribute to the pathogenesis of autoimmune diseases [23-27], determination of whether adenosine has a similar effect on Th1 and Th17 pathogenic T cell responses is important. In this study we show that the adenosine effect on Th17 responses is fundamentally different than its effect on Th1 responses; while it inhibits Th1 responses, it enhances Th17 responses. Mechanistic studies showed that the enhancing effect of adenosine on Th17 responses is accomplished via a sum of effects on various other cellular responses important for T cell activation, including αβ T cells, DCs and regulatory T cells. Adenosine is an important co-stimulating molecule for γδ T cell activation and augmented γδ T cell activation leads to high Th17 responses [28-31]. Adenosine is also an important molecule modulating DC differentiation [32-35]. In addition, adenosine exposed DCs showed a greater stimulating effect on γδ T cell activation leading to enhanced Th17 responses. Adenosine and γδ-based treatments should be more successful if the mechanisms by which they affect Th1 and Th17 responses is better understood.

## Materials and Methods

### Animals and Reagents

All animal studies conformed to the Association for Research in Vision and Ophthalmology statement on the use of animals in Ophthalmic and Vision Research. Institutional approval by Institutional Animal Care and Use Committee (IACUC) of Doheny eye institute, University of Southern California was obtained and institutional guidelines regarding animal experimentation followed.

Female C57BL/6 (B6) and TCR-δ^-/-^ mice on the B6 background, purchased from Jackson Laboratory (Bar Harbor, ME). They were housed and maintained in the animal facilities of the University of California in Los Angeles (UCLA). Recombinant murine IL-1β, IL-7, and IL-23 were purchased from R & D (Minneapolis, MN). Fluorescein isothiocyanate (FITC)-, phycoerythrin (PE)-, or allophycocyanin (APC)-conjugated antibodies (Abs) against mouse CD4, αβ T cell receptor (TCR), or γδ TCR and their isotype control antibodies were purchased from Biolegend (San Diego, CA). (PE)-conjugated anti-mouse IFN-γ and IL-17 monoclonal antibody was purchased from Santa Cruz Biotechnology (Dallas, Texas). The non-selective AR agonist 50-N-ethylcarboxamidoadenosine (NECA), selective A2AR agonist 2-p-(2-carboxyethyl) phenethylamino-5’-N-ethylcarboxamidoadenosine (CGS21680), selective A2AR antagonist (SCH 58261) [36, 37], were purchased from Sigma-Aldrich (St. Louis, MO, USA). Toll ligands were purchased from Invivogen (San Diego, CA).

### T cell preparations

αβ T cells were purified from B6 mice immunized with the human interphotoreceptor retinoid-binding protein (IRBP) peptide IRBP_1-20_, as described previously [28-30], while γδ T cells were purified from immunized and control (naïve) B6 mice. Nylon wool-enriched splenic T cells from naive or immunized mice were incubated sequentially for 10 min at 4°C with FITC-conjugated anti-mouse γδ TCR or αβ TCR Abs and 15 min at 4°C with anti-FITC Microbeads (Miltenyi Biotec GmbH, Bergisch Gladbach, Germany), then the cells were separated into bound and non-bound fractions on an autoMACS™ separator column (Miltenyi Biotec GmbH). The purity of the isolated cells, determined by flow cytometric analysis using PE-conjugated Abs against αβ or γδ T cells, was >95%. Resting γδ T cells were prepared either by isolation from naïve mice or by incubating activated γδ T cells in cytokine-free medium for 5-7 days, at which time they show down-regulation of CD69 expression [38]. Highly activated γδ T cells were prepared by incubating resting γδ T cells for 2 days with Abs against the γδ TCR (GL3) and CD28 (both 2 µg/ml, both from Bio-Legend, San Diego, CA), or cytokines combination (IL-1β,IL-7 and IL-23).

### Measurement of Th1 and Th17 Responses

αβ T cells (1.8 x 10^6^) were collected from IRBP_1-20_-immunized B6 mice, with or without oxATP treatment, on day 13 post-immunization. To obtain a sufficient number of cells, we routinely pool the cells obtained from all six mice in the same group, before the T cells are further enriched. The cells were co-cultured for 48 h with irradiated spleen cells (1.5 x 10^6^/well) as antigen presenting cells (APCs) and IRBP_1-20_ (10 μg/ml) in a 24-well plate under either Th1 (culture medium supplemented with 10 ng/ml of IL-12) or Th17 polarized conditions (culture medium supplemented with 10 ng/ml of IL-23) [30, 39]. Cytokine (IFN-γ and IL-17) levels in the serum and 48 h of culture supernatants were measured by ELISA (R & D Systems). The percentage of IFN-γ^+^ and IL-17^+^ T cells among the responder T cells was determined by intracellular staining 5 days post in vitro stimulation, and followed by FACS analysis, as described previously [39].

### Generation of Bone Marrow Dendritic Cells

Bone marrow dendritic cells (BMDCs) were generated by incubation of bone marrow cells for 5 days in the presence of 10 ng/ml of recombinant murine GM-CSF and IL-4 (R&D Systems), as described previously [40]. Cytokine (IL-1β, IL-6, L-12 and IL-23) levels in the culture medium were measured by ELISA. To determine antigen-presenting function, BMDCs were incubated in a 24-well plate with responder T cells isolated from immunized B6 mice under Th1-or Th17-polarizing conditions. Forty-eight hours after stimulation, IFN-γ and IL-17 in the culture medium were measured by ELISA. The percentage of IFN-γ^+^ and IL-17^+^ T cells among the responder T cells was determined by intracellular staining after 5 days of culture as described above.

### CFSE assay

Purified αβ T cells from IRBP1-20-immunized B6 mice were stained with CFSE (Sigma-Aldrich) as described previously [34]. Briefly, the cells were washed and suspended as 50 x 10^6^ cells/ml in serum-free RPMI 1640 medium (Corning Cellgro, VA), then were incubated at 37°C for 10 min with gentle shaking with a final concentration of 5 μM CFSE before being washed twice with RPMI 1640 medium containing 10% fetal calf serum (Atlantic Inc. Santa Fe, CA) (complete medium), suspended in complete medium, stimulated with immunizing peptide in the presence of irradiated syngeneic spleen cells as antigen-presenting cells (APCs), and analyzed by flow cytometry.

### Cytokine assays

Purified αβ T cells (3×10^4^ cells/well; 200 μl) from the draining lymph nodes and spleens of IRBP_1-20_-immunized B6 mice were cultured in complete medium at 37° C for 48 h in 96-well microtiter plates with irradiated syngeneic spleen APCs (1×10^5^) in the presence of 10 μg/ml of IRBP_1-20_, then a fraction of the culture supernatant was assayed for IL-17 and IFN-γ using ELISA kits (R & D).

## Statistical analysis

The results in the figures are representative of one experiment, which was repeated 3-5 times. The statistical significance of differences between groups in a single experimental was initially analyzed by ANOVA, and if statistical significance was detected the Student– Newman–Keuls post-hoc test was subsequently used. P values less than 0.05 was considered a statistically significant difference and marked with ** when P<0.05.

## Results

### Adenosine preferentially inhibits Th1 but not Th17 responses

To determine the adenosine effect on Th1 and Th17 responses in EAU, CD3^+^ responder T cells were harvested 13 days post immunization from the spleens and draining lymph nodes of B6 mice immunized with an uveitogenic antigen (IRBP_1-20_). The responder T cells were stimulated in vitro with the immunizing peptide and APCs (irradiated spleen cells), in the absence or presence of a selective A2AR agonist [2-p-(2-carboxyethyl) phenethylamino-5’-N-ethylcarboxamido-adenosine (CGS21680)], under culture conditions that favor Th17 or Th1 autoreactive T cell expansion (medium containing 10 ng/ml, respectively, IL-23 or IL-12) [30, 41]. Th1 and Th17 responses specific for the immunizing antigen were estimated by assessing responding IFN-γ^+^ and IL-17^+^ T cells after intracellular staining with fluoresce-labeled anti-IFN-γ or anti–IL-17 antibodies (Fig/1A). The results showed that the number of IFN-γ^+^ cells in response to CGS21680) decreased significantly, whereas the number of IL-17^+^ T cells remained unchanged. Alternatively, we used a CFSE assay (Fig.1B), in which the responder cells were pre-labeled with CFSE before stimulation under polarizing conditions. The results show that Th1 responses were readily inhibited by a very low dose (20 nM) of the A2AR agonist at a low dose that is inhibitory for Th1 response; but the Th17 responses remained minimally affected unless a very high dose (> 200nM) of the A2AR agonist was tested. Measurement of cytokine production of the responder T cells showed that IFN-γ production was inhibited by very low dose (20 nM) of A2AR agonist while IL17 production was only inhibited by 10 times higher doses of A2AR agonist (Fig.1C).

### γδ T cells offset an inhibitory effect of A2AR agonist on Th17 responses

Previous studies showed that γδ T cells are important enhancers of Th17 responses [38, 42, 43]. To determine the mechanism by which A2AR agonist is more inhibitory for Th1 responses than Th17 response, we compared Th17 responses in the presence or absence of γδ T cells. The CD3^+^ T cells containing γδ T cells were purified from immunized B6 mice and those not containing γδ T cells were prepared from immunized TCR-δ^-/-^ mice. The T cells were stimulated in vitro with the immunizing peptide and APCs, and the Th1 and Th17 responses determined by the number of αβTCR^+^IFN-γ^+^ and αβTCR^+^IL-17^+^ T cells among responder T cells and the amount of IFN-γ and IL-17 produced in culture supernatants by ELISA. The results in Fig. 2A showed that the generation of αβTCR^+^IL-17^+^ cells from wild-type (WT) B6 responders was enhanced by the A2AR agonist CGS21680 (Fig.2A, top panels) but not from TCR-δ^-/-^ responders (Fig2A, middle panels); moreover, if 2% of γδ T cells were added to TCR-δ^-/-^ responder T cells before in vitro stimulation their responses were also enhanced (Fig.2A lower panels) suggesting that γδ T cells in responder T cells counteracted any inhibitory effect of adenosine leading to greater Th17 responses. When IFN-γ- and IL-17-production were determined after in vitro stimulation (Fig.2B) in the presence of various agonists specific for different adenosine receptors, IL-17 production was inhibited by the CGS21680 but only in TCR-δ^-/-^ responders (Fig2B, top panels), not in B6 responder, indicating that inhibition was only seen in responder T cells lacking γδ T cells. The IFN-γ production of both responders were inhibited regardless whether γδ T cells were absent or present (Fig.2B, lower panels, indicating that Th1 inhibition was not dependent on γδ T cells. Studies comparing the effect of agonists specific for different adenosine receptors A1R, A2AR and A2BR, showed that agonists for A2BR and A1R adenosine receptors were also ineffective in inhibiting IL-17 production (Fig.2B). Since A2ARs are not strictly expressed on γδ T cells, we also compared the adenosine effect on Th17 responses of TCR-δ^-/-^ responder T cells supplemented with A2AR^+/+^ (from B6 mice) or A2AR^-/-^ γδ T cells (from A2AR^-/-^ mice). The results (Fig.2C) showed that adenosine was unable to enhance the Th17 responses supplemented with A2AR^-/-^ γδ T cells, indicating that binding of A2ARs to γδ T cells crucially involved adenosine-enhanced Th17 responses.

**Figure.1.**
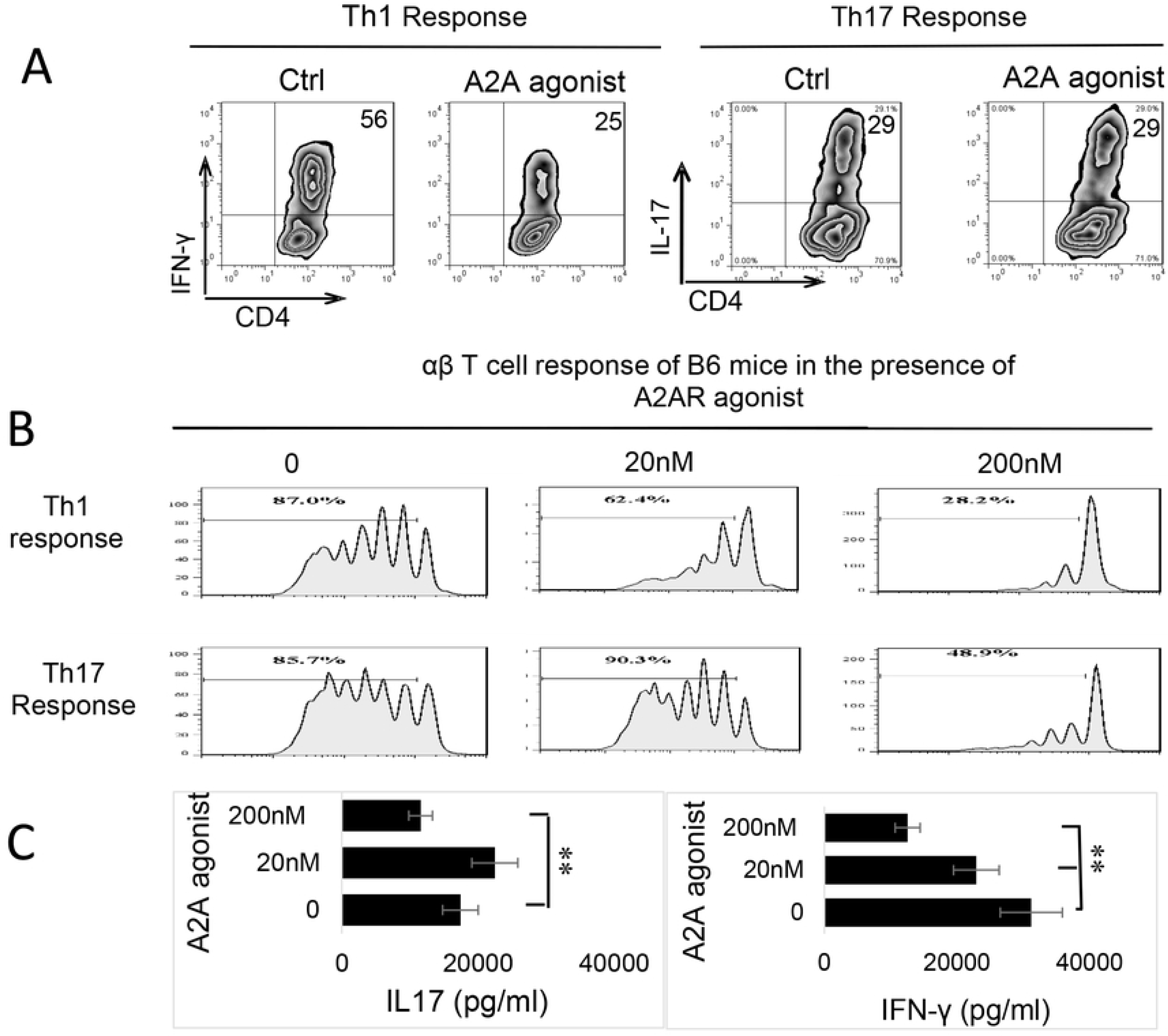
Adenosine’ effect on Th17 responses differed from its effect on Th1 responses. A) B6 mice were immunized with IRBP_1-20_/CFA. 13 days after immunization, CD3^+^ cells were separated from spleen and draining lymph nodes cells using MACS column. They were stimulated with the immunizing peptide (IRBP_1-20_) and APCs, in the absence or presence of an A2AR agonist (CGS21680), under Th17 (right panels) or Th1 (left panels) polarized conditions. The numbers of αβTCR^+^ IL-17^+^ and αβTCR^+^ IFN-γ^+^ cells were assessed after a 5-day in vitro stimulation by FACS analysis. B) CFSE assay for assessing dose-dependent effect (0-200 nM) of A2AR agonist (CGS21680) on Th1 and Th17 response. MACS column-separated CD3^+^ cells of immunized B6 mice were stimulated with the immunizing peptide (IRBP_1-20_) and APCs, under Th17 or Th1 polarized conditions, in the presence of indicated doses of CGS21680. The numbers of activated T cells were assessed by FACS analysis after a 5-day in vitro stimulation. The results shown are representative of those from five experiments. C) Calculated inhibition of Th1 and Th17 response by graded doses of CGS21680. **p < 0.05.

**Figure 2.**
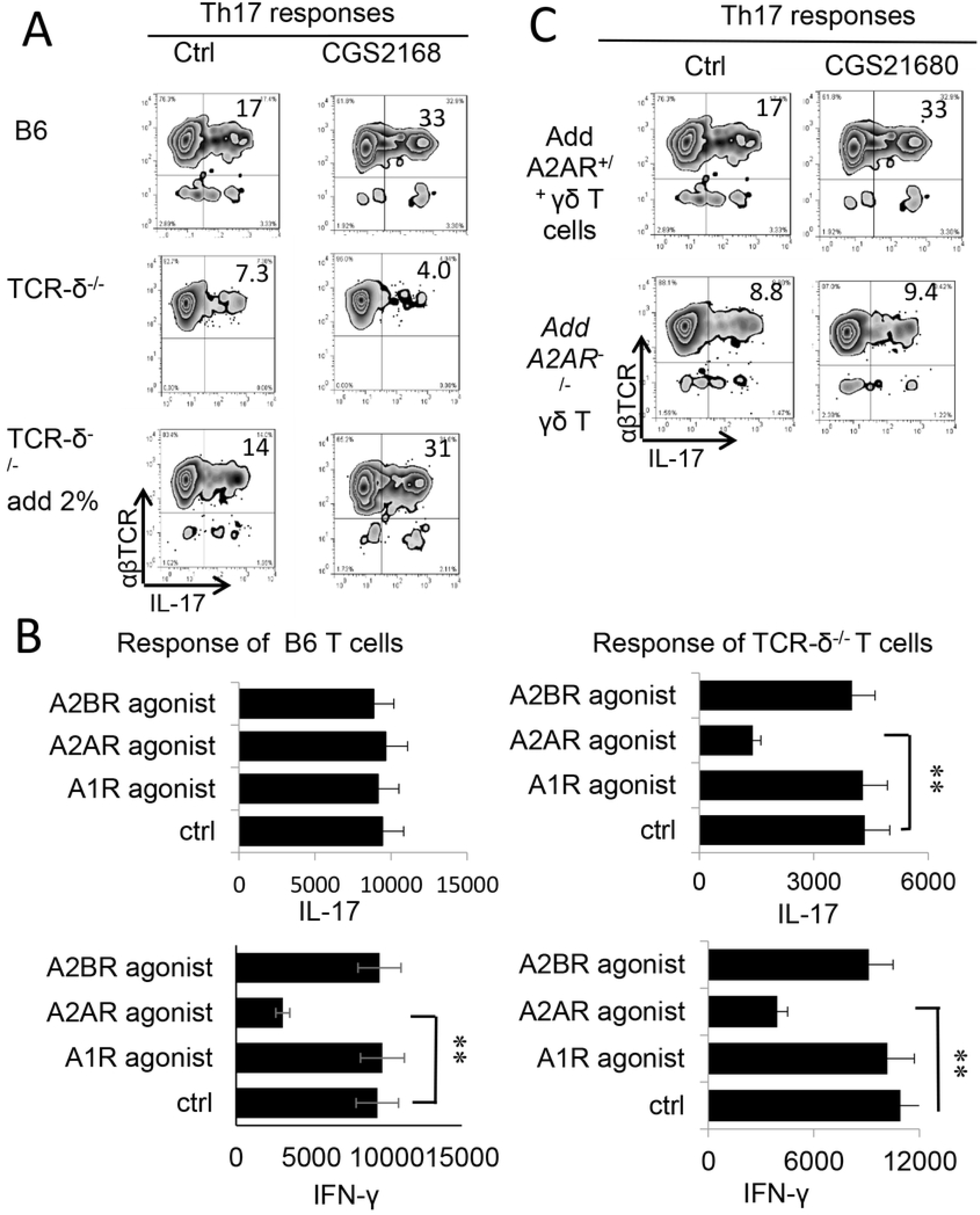
γδ T cell offsets the inhibitory effect of A2AR agonist on Th17 responses. A) Intracellular staining of IFN-γ^+^ and IL-17^+^ T cells among the responder T cells. Responder T cells were separated from either immunized B6 (top panels) TCR-δ^-/-^ mice without (mid panels) or with (lower panels) 2% supplemented γδ T cells. After stimulated with the immunizing peptide (IRBP_1-20_) and APCs, under Th17 polarized conditions. The numbers of αβTCR^+^ IL-17^+^ and αβTCR^+^ IFN-γ^+^ cells were assessed by FACS analysis after a 5-day in vitro stimulation. B) ELISA test assesses IL-17 (upper panels) and IFN-γ production (lower panels) by B6 (left panels) and TCR-δ^-/-^ responder T cells (right panels) under effect of agonists for specific adenosine receptors A1R, A2AR and A2BR. **, p < 0.05. C) Th17 responses of TCR-δ^-/-^ responder T cells were enhanced by A2AR^+/+^, but not A2AR^-/-^ γδ T cells. Responder T cells of TCR-δ^-/-^ mice were supplemented by 2% A2AR^+/+^ or A2AR^-/-^ γδ T cells, before stimulating with the immunizing peptide (IRBP_1-20_) and APCs, under Th17 polarized conditions.

### Adenosine augmented the Th17, but not Th1-stimulating effect of BMDCs triggered by a TLR ligand

DCs are principal antigen-presenting (AP) cells for initiating immune responses. Previous studies showed that TLR ligands have a profound effect on DC differentiation and maturation [44]. Since the levels of extracellular adenosine increases greatly during inflammation [11, 45, 46], we questioned whether adenosine and TLR ligands have counteractive or synergistic effects on DC function and Th1 and Th17 responses. To do so, we assessed GM-CSF-cultured BMDCs for AP effect in Th1 and Th17 responses, before and after exposure to adenosine and/or TLR ligands. The responder T cells were co-cultured with the treated BMDCs at ratio of DC:T=1:10 in the presence of immunizing antigen and measured cytokine production of responder T cells. After BMDCs were treated with LPS only, both IFN-γ and IL-17 production were increased. Unexpectedly, when BMDCs were treated with LPS and A2BR agonists IFN-γ and IL-17 production changed in opposite directions – IL-17 increased whereas IFN-γ declined (Fig.3B). Thus, BMDC Th1 and Th17-stimulating effects were dissociated under a dual effect of TLR ligand and adenosine, tipping the Th1 and Th17 balance towards the latter. We then investigated whether the higher Th17-promoting effect of adenosine was associated with altered cytokine production by BMDCs after exposure to LPS and/or adenosine. Our results showed that BMDCs did not produce the cytokines tested before the LPS exposure (not shown); treatment with either LPS (TLR4 ligand) or PAM3 (TLR2 ligand) stimulated a low production of all tested cytokines, including IL-12, IL-23, L-1β and IL-6. After LPS and adenosine, IL-12 production declined, and IL-23 production further increased indicating the dissociated Th1 and Th17 responses has been partly contributed by altered cytokine production of BMDCs. Given that IL-23 [25, 47] and IL-1β [48-50] has a strong Th17-promoting effect, changes in patters and amounts of cytokine production by BMDCs after adenosine presumably contributed to enhanced Th17 T cell response (Fig.3C).

**Figure 3.**
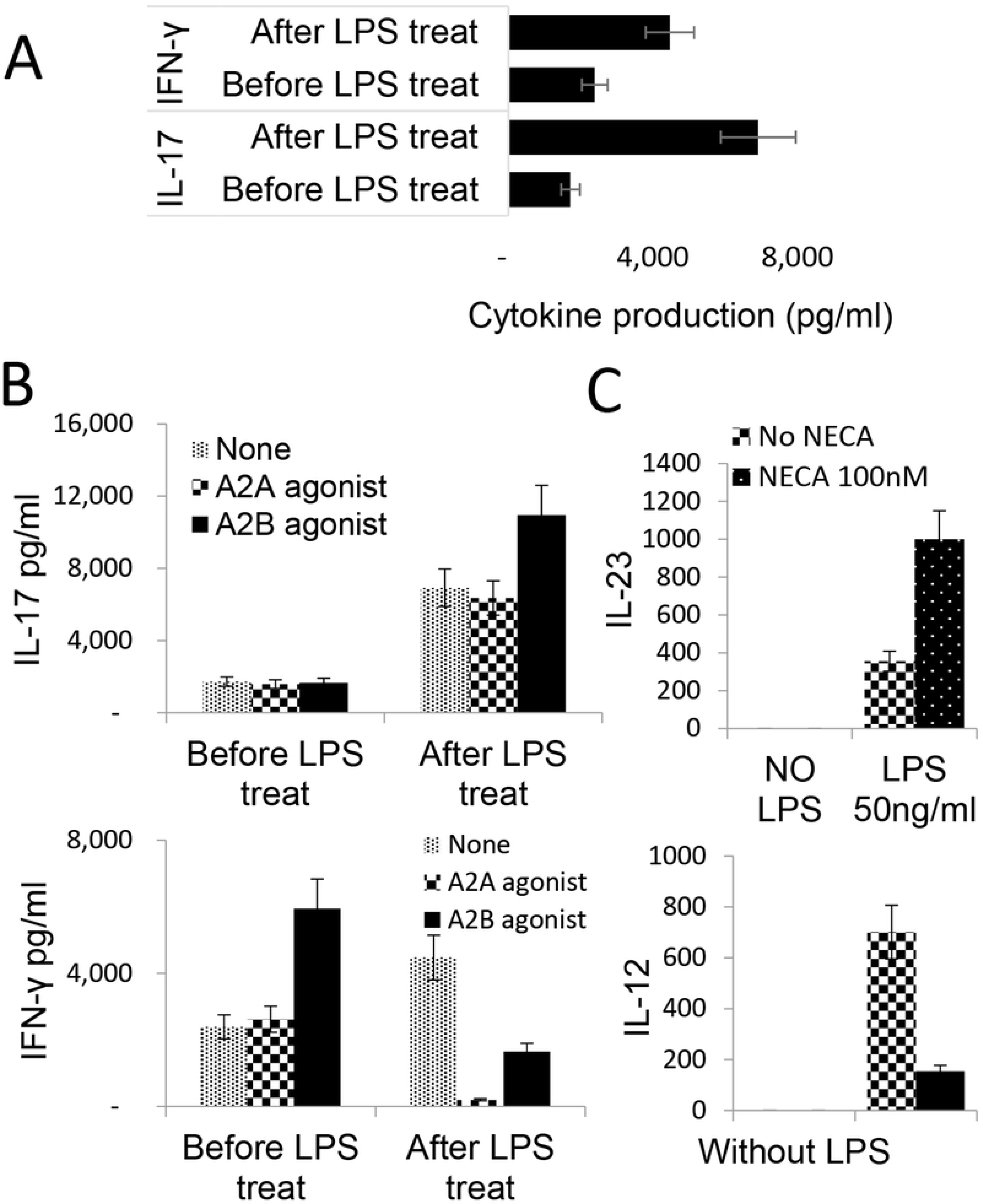
Adenosine augmented the Th17-, but not Th1-, stimulating effect of BMDCs triggered by TLR ligand. A) LPS treated BMDCs acquired increased stimulating effect on Th1 and Th17 responses. Responder T cells were isolated from immunized B6 mice. They were stimulated with the immunizing peptide (IRBP_1-20_) and BMDCs, under Th1 (upper panels) or Th17 (lower panels) polarized conditions. Cytokines in the supernatants were assessed by ELISA 48hr after stimulation. **, p < 0.05. B) Dissociated Th1 and Th17 stimulating effect of BMDCs after dual treatment with LPS and adenosine. BMDCs were treated with A2AR agonist or A2BR agonist (100 nM) before (left panels) or after (right panels) LPS treatment. After co-cultured with responder T cells, IFN-γ and IL-17 amounts in culture supernatants were determined by ELISA. The results show that after LPS treatment, A2BR agonist treatment augmented BMDCs’ Th17-stimulating effect, whereas both A2AR and A2BR agonists decreased BMDCs’ Th1-stimulating effect. C) IL-12 and IL-23 production by BMDCs after treated with LPS, with or without AR agonist. BMDCs produce IL-12 and IL-23 only after treated with LPS. When LPS treated BMDCs further exposed to AR agonist, the IL-12 production was declined, whereas the IL-23 production significantly increased.

### Adenosine augmented cytokine-mediated γδ T cell activation

Given our previous findings that γδ T cell activation was a major contributor in regulation of Th17 responses, we questioned whether the enhancing effect of adenosine on Th17 responses was due to augmented γδ T cell activation. As we have previously reported, purified γδ T cells can be activated by a number of proinflammatory cytokines and that a mixture of IL-1β, IL-7, and IL-23 has a strong stimulatory effect [30]. We used this combination and tested the activation of γδ T cells by cytokines and in the absence or presence of adenosine. Responder γδ T cells were prepared from immunized B6 mice using MACS sorter. Fig.4A showed that cytokines IL-1β, IL-7, and IL-23 were able to activate IL-17 production of γδ T cells; furthermore, a combination of adenosine analogue NECA and the cytokine mixture greatly augmented IL-17 production by γδ activation, even though neither NECA nor A2AR agonist itself did not appreciably stimulate γδ T cells. A similar synergistic effect was seen when γδ T cells were exposed to a combination of a TLR ligand and NECA (not shown). Assessment of in vivo effect of adenosine on γδ T cells showed that B6 mice that received an A2AR agonist (BAY60-6538) injection after immunized have greater numbers of γδ T cells among which the CD44^high^ γδTCR^+^ cells are more abundant Fig.4B. To further determine that adenosine is responsible for γδ T cell activation we also compared the activation of A2AR^+/+^ and A2AR^-/-^ γδ T cells by these cytokines. Our results showed that after stimulation with the same dose of cytokines the activation of A2AR^-/-^ γδ T cells was significantly lower compared to A2AR^+/+^ γδ T cells because the adenosine receptor A2AR^-/-^ on γδ T cells was disabled (Fig.4C).

**Figure 4.**
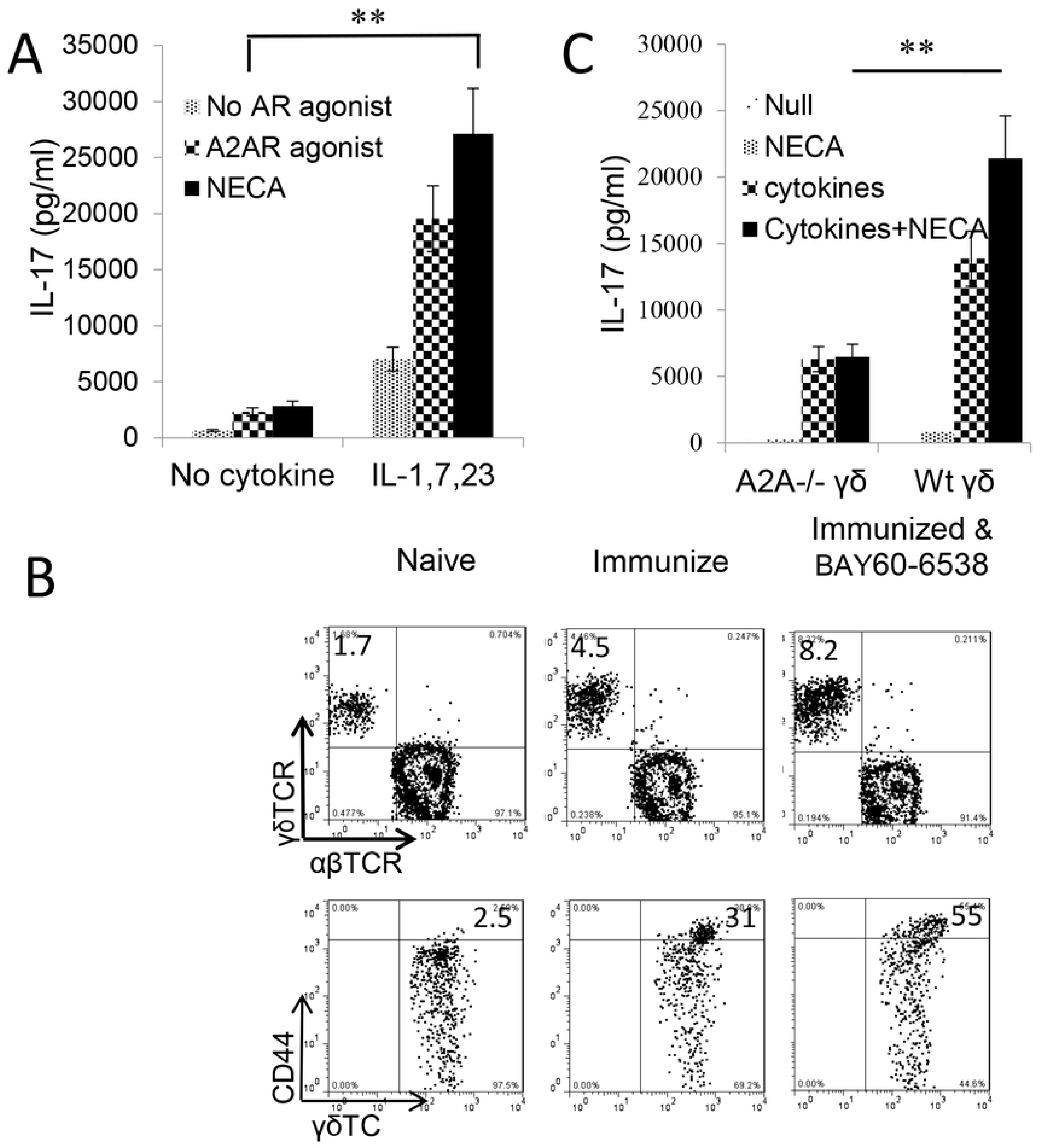
Adenosine analogue NECA augmented cytokine-mediated γδ T cell activation. A) IL-17 production by γδ T cells stimulated by cytokines is augmented by adenosine analogue NECA. MACS purified γδ T cells were isolated from immunized B6 mice. They were exposed to cytokines (IL-1β+7+23), in the absence or presence A2AR/A2BR agonists. IFN-γ and IL-17 in the cultured cell supernatants were assessed by ELISA. Results show that after, but not before, cytokine exposure, γδ T cells acquired increased response to A2AR agonist and NECA. **, p < 0.05. B) The number (upper panels) and activation status (lower panels) of γδ T cells are increased in immunized B6 mice administered with A2BR agonist (Bay60-6538). B6 mice were immunized with IRBP_1-20_/CFA with or without an injection of A2BR agonist (Bay60-6538). 13 days pos-immunization CD3^+^ cells isolated were assessed for abundance as well as activation status of γδ T cells. The gated CD3^+^ cells (upper panels) and γδTCR^+^ T cells (lower panels) were analyzed. C) γδ activation is compromised if A2ARs on γδ T cells are disabled. γδ T cells isolated from B6 (A2AR^+/+^) or A2AR^-/-^ mouse (A2AR^-^/^-^) were compared for response to cytokines (IL-1β+7+23) and/or NECA. γδ activation is accessed by measurement of IL-17 production. Results show that Adenosine enhances A2AR^+/+^ but not A2AR^-^/^-^ γδ activation. The results shown are from a single experiment and are representative of those obtained in >5 experiments.

### Adenosine augmented the TLR ligand activation of γδ T cells by BMDCs

An alternative pathway of γδ T cell activation is stimulation by DCs. To determine whether BMDCs exposed to adenosine acquired an increased ability to stimulate γδ T cells, GM-CSF cultured BMDCs were co-incubated with MACS-sorted γδ T cells, after treatment with LPS and/or NECA, at a ratio of T:DC=10:1 for two days. The activation of γδ T cells was assessed by measuring IL-17 production and the numbers of CD69^+^ γδ T cells. The results showed that (Fig.5A) only the only LPS treated BMDCs could stimulate γδ T cells to produce IL-17 and BMDCs treated with LPS plus NECA acquired a greater stimulating effect (Fig.5A). However, BMDCs treated with NECA alone were not stimulatory, indicating that the effect of adenosine on BMDCs is indirect and needed to be synergized with cytokines. Expression of CD69 – a cell surface marker identifying activated T cells - showed that only activated γδ T cells stimulated by LPS treated BMDCs could augment γδ activation leading to augmented Th17 responses; furthermore, treatment of BMDCs with LPS plus NECA further augmented the stimulating effect (Fig.5B) of adenosine.

**Figure 5.**
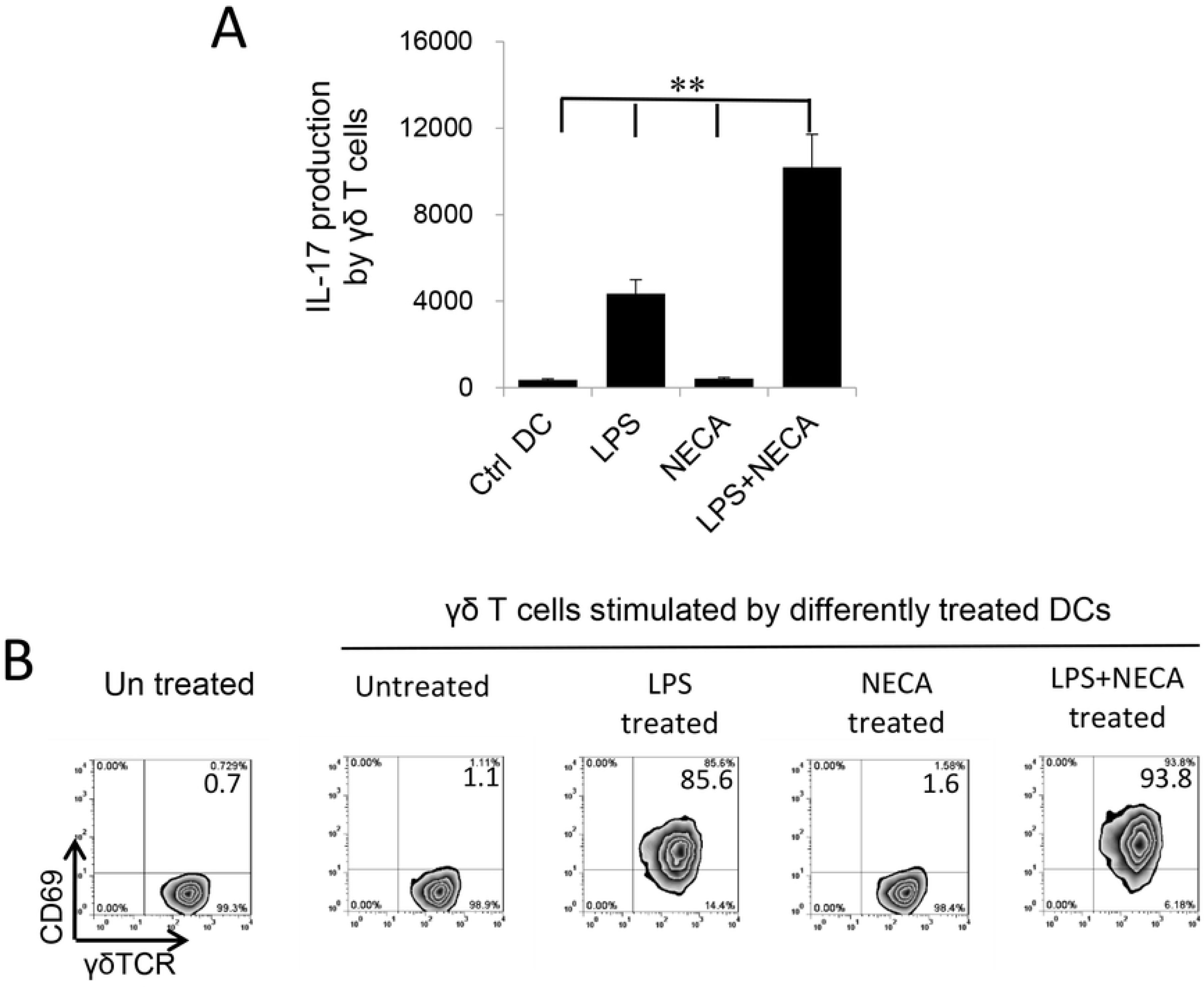
Synergistic effect of NECA and TLR ligand in BMDCs’ γδ-stimulating activity. A) BMDCs-stimulated γδ T cells produced increased amounts of IL-17 if BMDCs were pre-treated with LPS and/or NECA. B) BMDCs-stimulated γδ T cells expressed increased levels of CD69, if BMDCs were pre-treated with LPS and/or NECA. The gated γδTCR^+^ T cells were further analyzed.

### Inhibition of ADA by an ADA inhibitor augmented the IL17 responses

Endogenously produced adenosine is degraded by amino deaminase (ADA). We observed that Toll ligands activated BMDCs expressed increased amounts of ADA (Fig6A). To determine whether Th1 and Th17 responses would be affected if ADA function is deactivated, we determined the AP function of BMDCs with or without prior treatment with EHNA [erythro-9-(2-hydroxy-3-nonyl)] - a reversible inhibitor of ADA ^[51, 52]^. The results showed that inhibition of ADA by EHNA enhanced the Th17 as well as γδ T cell responses (Fig.6B). Measurement of cytokine production of BMDCs showed that untreated BMDCs produced neither IL-23 nor IL-1β, but the production of these cytokines was induced by LPS. Adenosine inhibited LPS-induced cytokine production; however, the production increase significantly if simultaneously treated with EHNA. These results supported the prediction that regulation of endogenously generated adenosine by ADA crucially controls adenosine levels, and thus, Th17 responses; when ADA is disabled, adenosine will accumulate and Th17 responses will be enhanced.

**Fig. 6.**
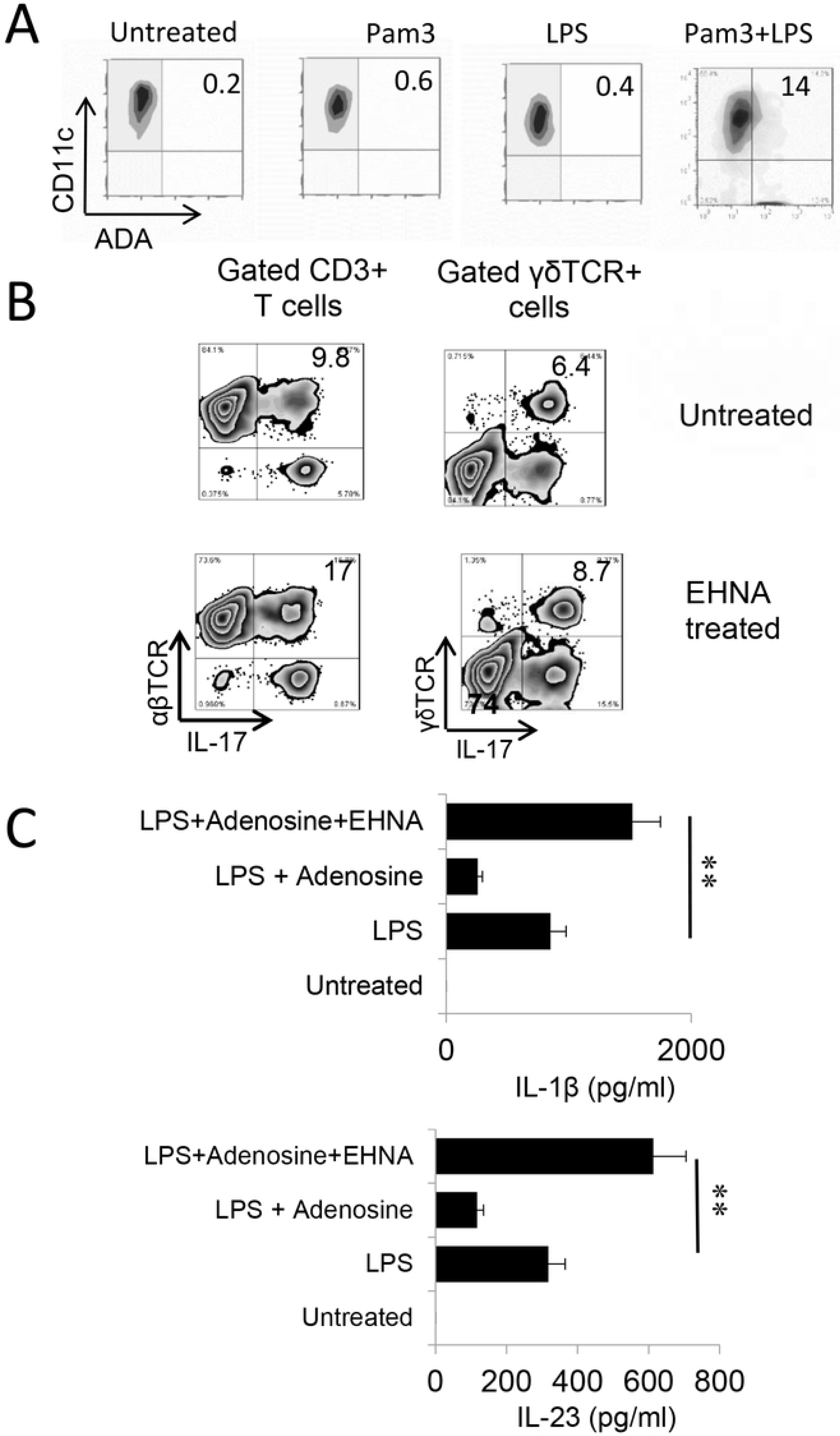
ADA inhibitor EHNA inhibited Th1 responses but enhanced the Th17 responses. A) BMDCs express increased levels of ADA after TLR ligands stimulation. BMDCs were incubated with LPS, Pam3, or LPS+Pam3 for 48h, before they were stained with an anti-ADA antibody. B) Prior treatment of BMDCs with EHNA augmented Th17 responses. Responder CD3^+^ T cells isolated from immunized B6 mice were stimulated, under Th17 polarized conditions, with BMDCs with (upper panels) or without (lower panels) a prior treatment with EHNA (10 µM). Expressing IL-17 of responder T cells and gated γδ T cells were evaluated by FACS analysis after intracellular stain. Results show that inhibition of ADA by EHNA on BMDCs enhanced Th17 responses. C) BMDCs produced increased amounts of IL-1β and IL-23 after treatment with EHNA. BMDCs were treated with LPS (100ng/ml), LPS plus adenosine 1 μg/ml), or LPS plus adenosine and EHNA (10 µM) as indicated. IL-1β and IL-23 amounts in supernatants were tested 48 h after stimulation. Results show that while adenosine inhibited LPS-induced cytokine production, additional treatment of EHNA increased cytokine production. **, p < 0.05.

## Discussion

During stress and tissue injury, ATP is released from the intracellular compartment into the extracellular space, where it is degraded to adenosine through a cascade of enzymatic reactions. Elevated amounts of adenosine are found in ischemia, inflammation and trauma [1, 53-55]. Degradation of ATP to adenosine involves ectonucleotidases including CD39 (nucleoside triphosphate diphosphorylase [NTPDase]) and CD73 (5′-ectonucleotidase [Ecto5′NTase]) [3, 12]. Produced adenosine is degraded by amino deaminase (ADA) [51, 52]. Adenosine is an important regulatory molecule since it modulates a wide range of physiological functions [4] including the immune response [1, 4-6] by acting on many types of immune cells, including T cells [56, 57], macrophages/DCs [13, 58], NK cells [59], neutrophils [54], platelets [60], and regulatory T cells [13, 14, 61].

Four types of adenosine receptors (ARs) have been defined, designated A1R, A2AR, A2BR, and A3R [1, 11]. The major functional receptors on T cells is A2AR [11, 62]. Previous studies have demonstrated that adenosine has a direct inhibitory effect on αβ T cells and macrophages/dendritic cells [13, 33-35, 58, 63, 64]. Treatment with adenosine reduced Th1 responses [65, 66] and activation of A2AR on T cells inhibited T-cell– mediated cytotoxicity, cytokine production [67] and T-cell proliferation [68, 69] [14]. Regulatory T cells exert their suppressive action through the production of adenosine [68, 70, 71]. Adenosine inhibits IL-12 production by DCs via which Th1 responses are inhibited [72]. The extrapolation of adenosine as inhibitory was mostly obtained from studies of Th1 immune responses, since Th17 responses were discovered only recently. Given that both Th1 and Th17 pathogenic T cells contribute to the pathogenesis of autoimmune diseases [23-27], determination of whether adenosine has a similar effect on Th1 and Th17 pathogenic T cells is important. In this study, we show that the adenosine effect on Th17 responses is enhancing. As a result, it tips the Th1 and Th17 balance toward the latter.

To determine the mechanisms by which Th17 responses differed from Th1 autoreactive T cells in response to adenosine, we used a well-established mouse model of uveitis – experimental autoimmune uveitis (EAU). We found that adenosine plays an important role in modulating the pathogenesis of EAU and that it exerts distinct effects on Th1 and Th17 autoimmune responses. Prominently, adenosine suppresses Th1 autoimmune responses, but enhances Th17 responses. The opposite effect of adenosine on Th1 and Th17 responses could certainly offset therapeutic attempts to regulate Th1 pathogenic reactions. As such, clarification of the conflicting effect of adenosine on Th1 and Th17 responses is of major importance.

The enhancing effects of adenosine on Th17 responses is accomplished via several pathways, among which γδ T cell activation is most important. We have previously reported that γδ T cell activation is a key pathogenic event leading to augmented Th17 responses and the development of EAU [30, 31, 38, 73]. We therefore investigated whether the dissociated effect of adenosine on Th1 and Th17 responses involves its effect on γδ activation. Our results showed that adenosine enhances γδ activation, which in turn augmented Th17 responses [30, 31, 38]. In the absence of γδ T cells, adenosine is inhibitory for both Th1 and Th17 responder T cells; however, adenosine inhibits only Th1, but not Th17, responses when as few as 2% γδ T cells were present. The Th17 enhancing effect of γδ T cells was abolished when A2ARs on γδ T cells is disabled (Fig.2) supporting the notion that adenosine effect on γδ T cells’ enhancement of Th17 responses. Adenosine also promotes DC differentiation into a unique subset that strongly stimulated Th17, but not Th1, responses. It also augmented the γδ-stimulating activity of BMDCs, by which Th17 responses are further promoted.

In the study of DCs’ we found that BMDCs have Th1-stimulating activity but very little Th17 stimulating capacity before adenosine treatment. After treatment with TLR ligands, both Th1 and Th17 stimulating effects of BMDCs were enhanced. Unexpectedly, when BMDCs were treated with both TLR ligand and adenosine, the Th1 and Th17-stimulating effects of BMDCs dissociated - while the Th1-stimulating function declined, Th17 stimulation increased and tipped the Th1/Th17 balance towards the latter. Since levels of extracellular adenosine increases greatly during inflammation, we predict that a unique environmental condition occurs during inflammation. Given that ATP may function as an endogenously generated TLR ligand [18, 74, 75], it is likely that the balance of ATP and its degrading adenosine metabolites plays an important role in the T cell response. To investigate the function of ATP degradation and deactivation of adenosine by adenosine deaminase enzyme we examined whether deactivation of ADA by a specific enzyme (EHNA) would result in excess adenosine and promote cascading Th17 responses. Our results demonstrated that ADA inhibition favors enhanced Th17 responses.

Alternative pathways may have been also involved in adenosine-induced enhancement of T cell responses. As we previously reported, activated γδ T cells express greatly increased amounts of high-affinity adenosine receptors (A2ARs) [31, 73], leading to a greatly increased adenosine binding capability [73] The preferential binding of adenosine by γδ T cells may lead to a re-distribution of adenosine among various immune cells, leading to diminished adenosine binding by αβ T cells, for example, which will also favor augmented αβ T cell responses [30, 31, 38, 73].

A better knowledge and understanding of the functional conversion of adenosine should facilitate adenosine-mediated immunotherapies. The cellular and molecular basis for enhancing and/or inhibiting the effects of ATP/adenosine remain to be further determined and the outcome of such study should improve current available therapies, including adenosine- and γδ T cell-based immunotherapies.

## Footnotes

This work was supported by NIH grants EY0022403 and EY018827 and grant from for Research to Prevent Blindness, NYC.

## Supporting Information

S1 File. The ARRIVE guidelines checklist

